## Supporting Figures for "Conformational and oligomeric states of SPOP from small-angle X-ray scattering and molecular dynamics simulations"

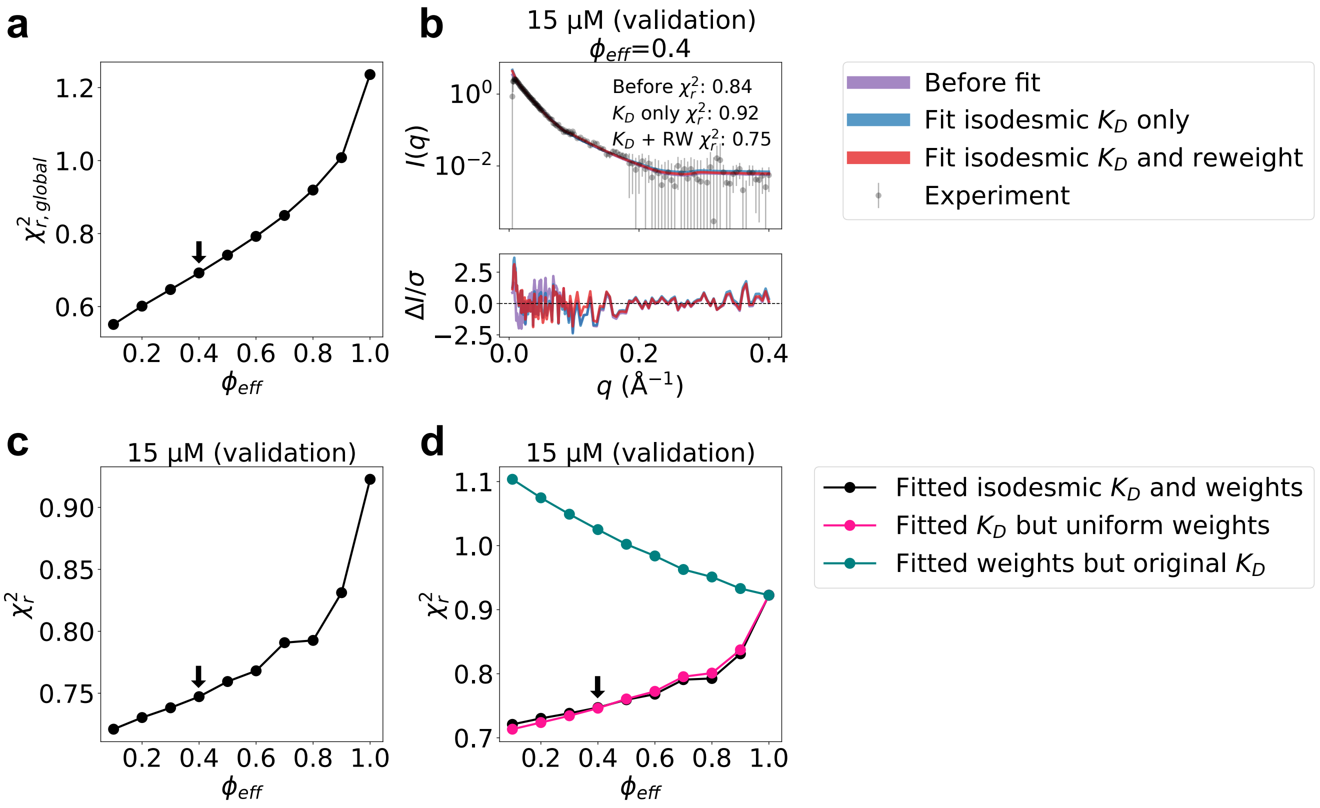


**Figure 1. Selection of $\phi_{\text{eff}}$ and model validation**. **a.** $\chi^2_{r,global}$ calculated from the concentration series of SAXS data as a function of the fraction of effective frames, $\phi_{\text{eff}}$, retained after BME reweighting. The arrow shows the selected value of $\phi_{\text{eff}}$. $\phi_{\text{eff}}$=1 corresponds to the MD simulations before reweighting. **b.** Agreement between calculated and experimental SAXS data recorded using 15 μM protein, which was not used for optimization. Calculated SAXS profiles are shown before fitting (isodesmic $K_D$=2.4 μM) (purple), with isodesmic $K_D$ optimized (isodesmic $K_D$=0.9 μM) (blue), and with isodesmic $K_D$ and ensemble weights optimized with $\phi_{\text{eff}}$=0.4 (isodesmic $K_D$=1.3 μM) (red). Error-normalized residuals are shown below the SAXS profile and $\chi_r^2$ for the three cases are shown on the plot. **c.** Validation using SAXS data at 15 μM protein. $\chi_r^2$ to the SAXS data at 15 μM, which was not used for optimization, using the ensemble weights and $K_D$ determined from the optimization as a function of $\phi_{\text{eff}}$. Only SAXS scale and constant background were fitted to the 15 μM SAXS data. The arrow shows the selected value of $\phi_{\text{eff}}$. **d.** Same as panel c (black), but also showing the agreement given by using only the fitted isodesmic $K_D$ with uniform weights (unbiased MD ensemble; red) or using only the fitted weights but the isodesmic $K_D$ of 0.9 μM determined before reweighting (green).

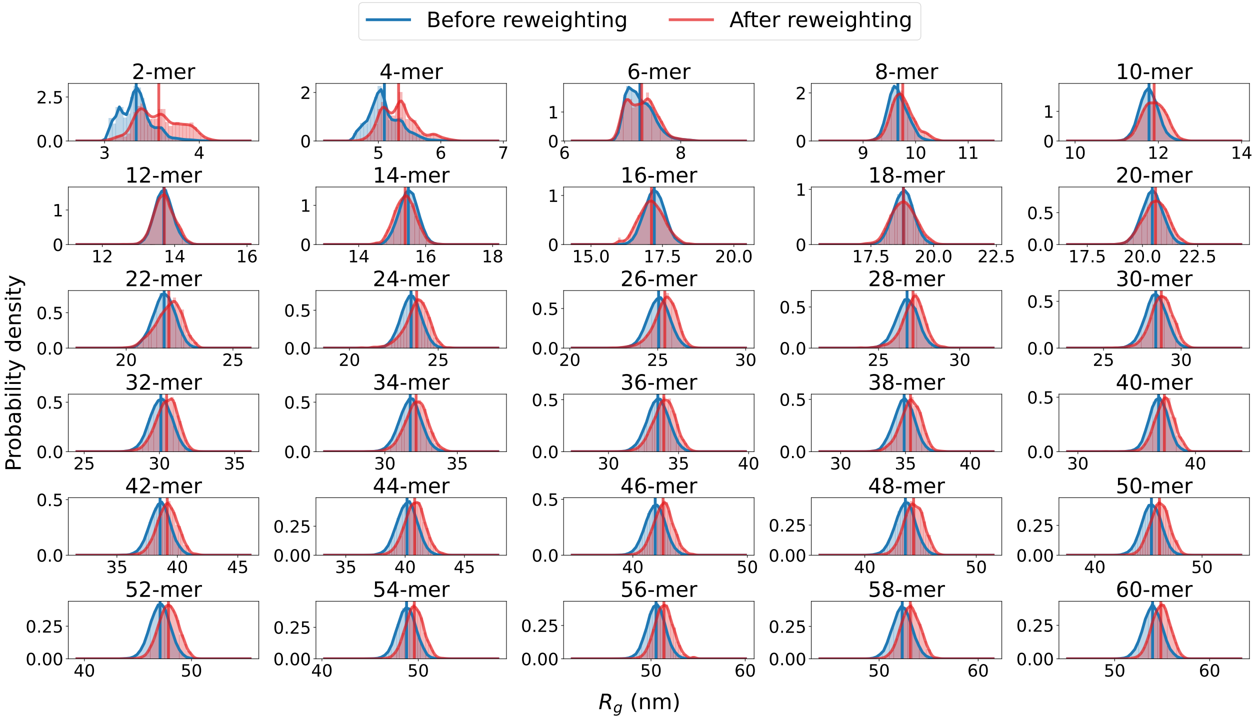


**Figure 2.** $R_g$ **distributions before and after reweighting**. Probability distribution of the radius of gyration ($R_g$), calculated from ensembles of SPOP oligomers before and after reweighting against SAXS data. Average values are shown as vertical lines.

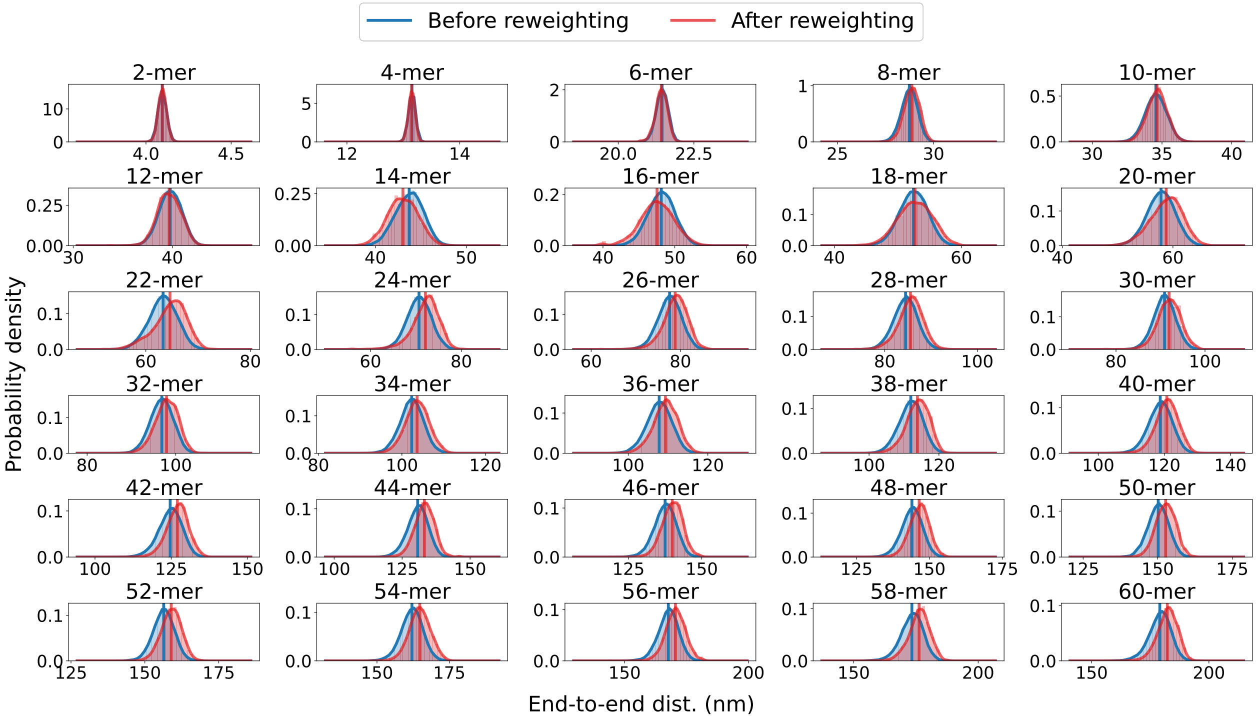


**Figure 3. End-to-end distance distributions before and after reweighting**. Probability distribution of the end-to-end distance, calculated from ensembles of SPOP oligomers before and after reweighting against SAXS data. Average values are shown as vertical lines.

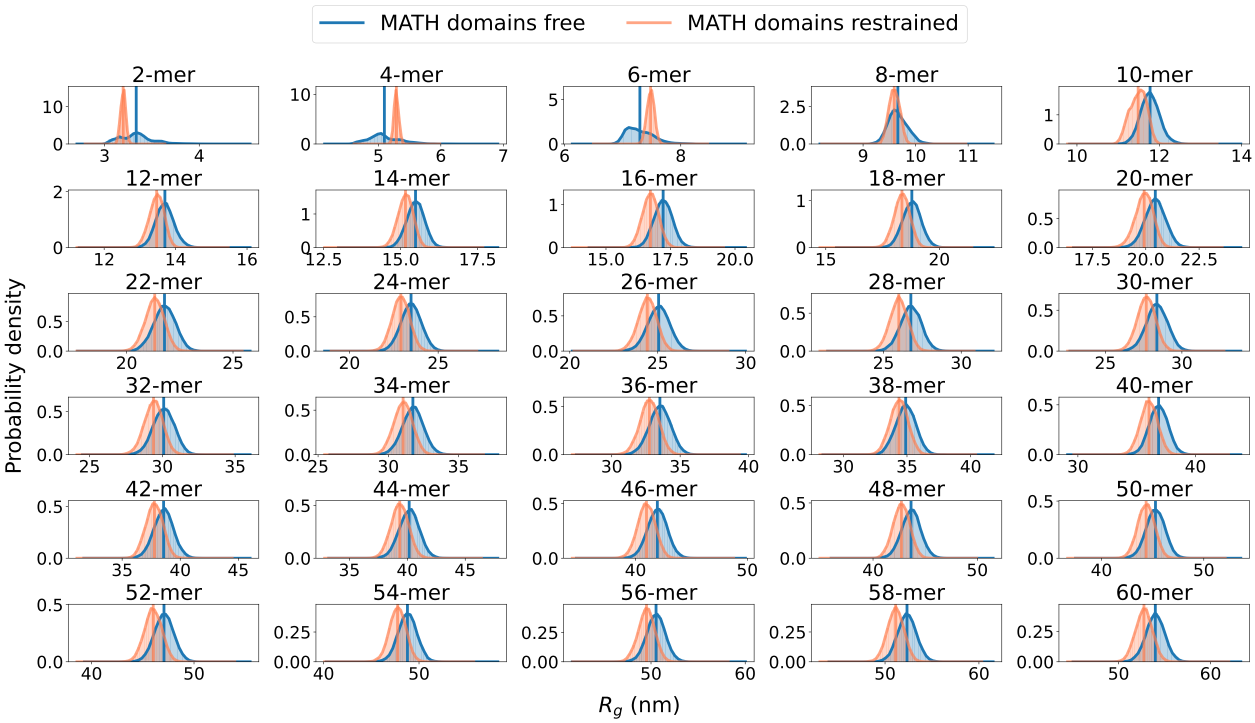


**Figure 4.** $R_g$ **distributions from simulations with MATH free and MATH restrained**. Probability distribution of the radius of gyration ($R_g$), calculated from ensembles of SPOP oligomers generated with MATH domains unrestrained (blue, free) or restrained to the BTB/BACK domains based on the configuration in the crystal structure using the Martini elastic network model (orange, restrained). Average values are shown as vertical lines.

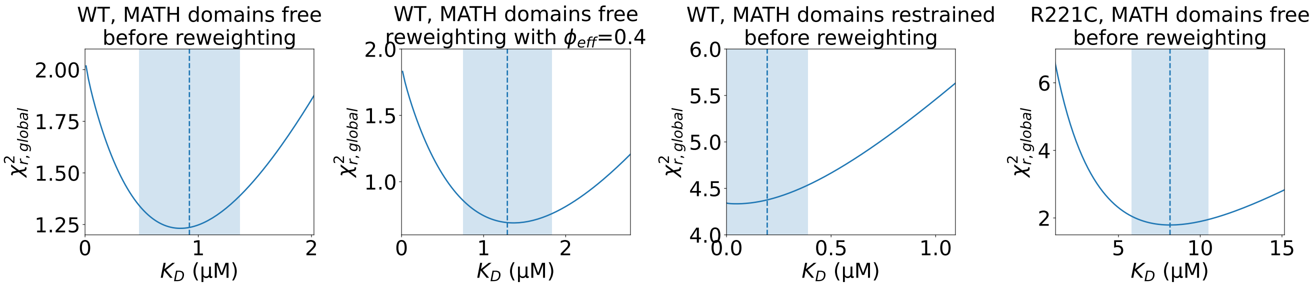


**Figure 5. Determining the error of the fitted isodesmic** $K_D$. $\chi^2_{r,global}$ to the concentration series of SAXS data given by the conformational ensembles shown above the plot with oligomer populations given by a range of isodesmic $K_D$ values around the $K_D$ fitted with simulated annealing. Only the SAXS scale and constant background were fitted for each $K_D$. The $K_D$ fitted with simulated annealing is shown as a dashed line and the selected error is shaded. The error was selected to include all $K_D$ values that give a $\chi^2_{r,global}$ no more than 10% greater than the minimum $\chi^2_{r,global}$.

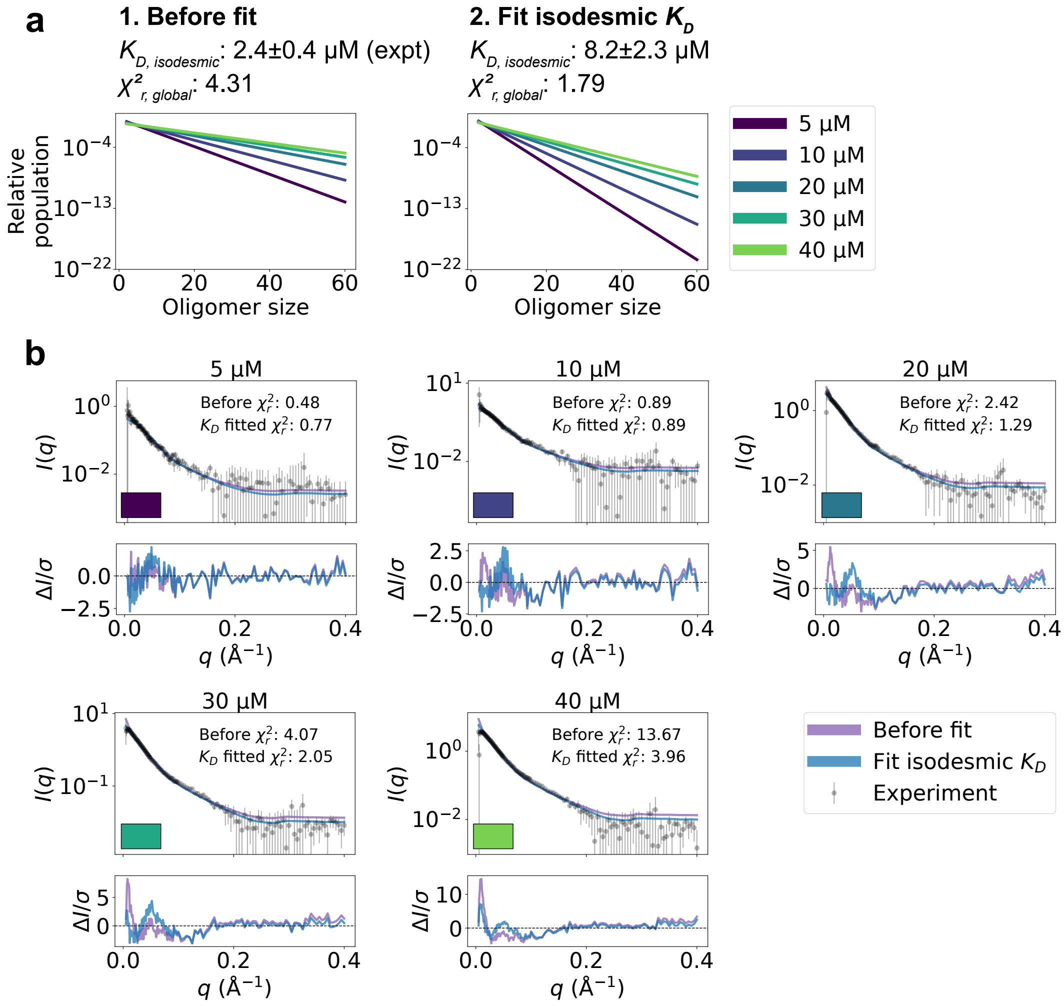


**Figure 6. Fit to SAXS data for SPOP R221C. a.** Relative populations of oligomers for the protein concentrations used in SAXS experiments (note the logarithmic scale). Populations are given by the isodesmic model with the $K_D$ value noted above the plot, which is either (1) previously determined with CG-MALS or (2) fitted globally to the SAXS data in panel b. $\chi^2_{r,global}$ quantifies the agreement with SAXS data in panel b for the two scenarios. **b.** Agreement between experimental SAXS data on SPOP R221C and averaged SAXS data calculated from conformational ensembles of SPOP oligomers with populations given by the isodesmic model (as shown in panel a). Error-normalized residuals are shown below the SAXS profiles and $\chi^2_r$ to each SAXS profile is shown on the plot.

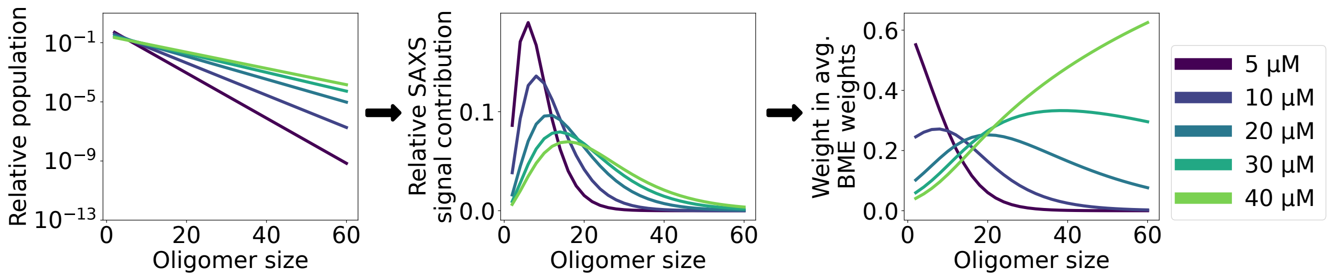


**Figure 7. Averaging the conformational weights from different SAXS experiments**. Left: relative oligomer populations given by the isodesmic model with fitted $K_D$=1.3 μM for each protein concentration in the SAXS concentration series. Middle: relative contribution of each oligomer to the averaged SAXS signal given the populations in left plot. Right: weight given to conformational weights obtained with each SAXS experiment when averaging to get a single set of conformational weights for each oligomer.
